# Morphogen-Lineage Selector Interactions During Surface Epithelial Commitment

**DOI:** 10.1101/348839

**Authors:** Sandra P. Melo, Jillian M. Pattison, Samantha N. Piekos, Jessica L. Torkelson, Elizaveta Bashkirova, Maxwell R. Mumbach, Charlotte Rajasingh, Hanson Hui Zhen, Lingjie Li, Eric Liaw, Daniel Alber, Adam J. Rubin, Gautam Shankar, Howard Y. Chang, Paul A. Khavari, Anthony E. Oro

## Abstract

Human embryonic stem cell (hESC) differentiation promises advances in regenerative medicine^1–3^, yet conversion of hESCs into tissues such as keratinocytes requires a better understanding of epigenetic interactions between the inductive morphogens retinoic acid (RA) and bone morphogenetic protein 4 (BMP), and the master regulator p63^4,5^. Here we develop a robust, defined, keratinocyte differentiation system, and use a multi-dimensional genomics approach to interrogate the contributions of the morphogens and lineage selector to chromatin dynamics during early surface ectoderm commitment. In stark contrast to other master regulators^6–9^, we find using p63 gain and loss of function hESC lines, that p63 effects major transcriptional changes only after morphogenetic action. Morphogens alter chromatin accessibility and histone modifications, establishing an epigenetic landscape for p63 to modify. In turn, p63 closes chromatin accessibility and promotes the accumulation of repressive H3K27me3 histone modifications at sites distal to where it binds. Surprisingly, cohesin HiChIP^10^ visualization of genome-wide chromosome conformation reveals that both p63 and the morphogens contribute to dynamic long-range genomic interactions that increase the probability of negative transcriptional regulation at p63 target loci. p63-regulated accessibility, not H3K27me3 deposition, appears to drive early transcriptional changes. We illustrate morphogen-selector interactions by studying p63 negative feedback regulation of *TFAP2Ci*^11^, whereby disruption of the single p63 binding site results in a loss of p63-mediated transcriptional control and dramatic increases in TFAP2C and p63 expression. Our study reveals the unexpected dependency of p63 on morphogenetic signaling to control long-range chromatin interactions during tissue specification and provides novel insights into how master regulators specify diverse morphological outcomes.

---

Published protocols of hESC-derived keratinocytes suffer from heterogeneity due to feeders and additive variability^5,12–15^, thus we developed a xeno-free, chemically-defined differentiation system based on E6 media^16^ supplemented with two morphogens, RA and BMP4 (Fig. 1a). This system was highly reproducible using hESCs and recapitulated commitment towards a surface ectoderm fate, indicated by immunofluorescence (IF) analysis of epithelial markers keratin 18 (K18)^17^ and p63^18,19^ by day 7, followed by high levels of p63 and the keratinocyte maturation marker keratin 14 (K14)^20^ by day 45 (Fig. 1a). Robust p63 expression occurred only when both morphogens were present, indicating a synergistic role for p63 accumulation (Fig. 1b,c, Extended Data Fig. 1). As morphogenetic exposure for 7 days induced both uniform p63 expression and subsequent keratinocyte development^4,5^, we interrogated this key 7-day stage with a multi-dimensional genomics approach to understand the functional interaction between p63 and the morphogens.

**Figure 1.**
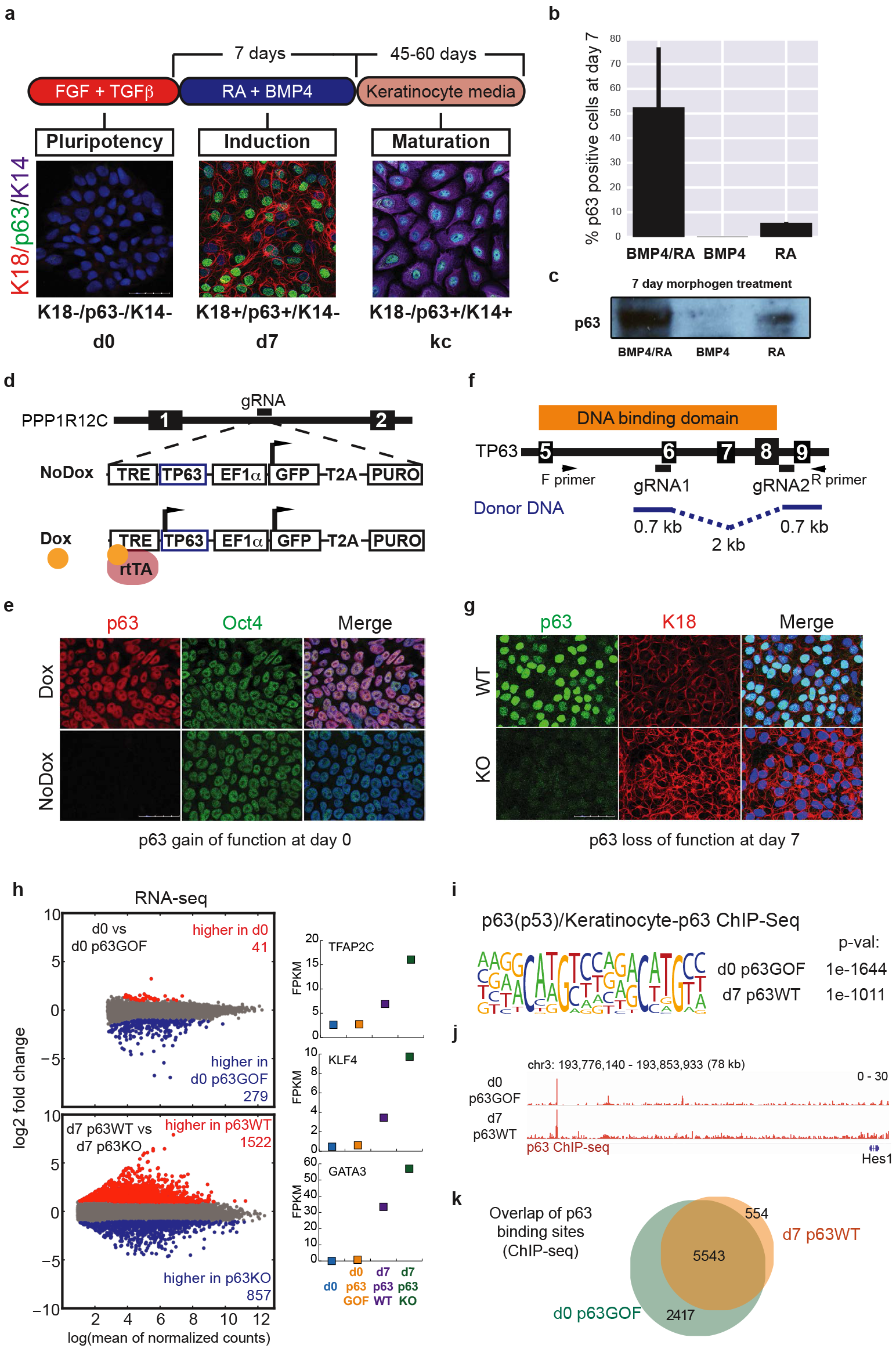
Morphogens and the lineage selector p63 cooperate to drive early stratified epithelial differentiation. (a) Differentiation of hESCs into keratinocytes takes 60 days in the xeno-free, defined system. Treatment with RA and BMP4 for 7 days induces K18 and p63 expression. Switching the cells into keratinocyte media allows for selection and growth of functional keratinocytes (kc) that express K14 and p63. (b,c) hESCs need exposure to both RA and BMP4 to achieve high p63 expression. Error bars represent standard deviation. (d) Strategy for generating the d0 p63GOF cell line. Numbered black boxes signify exons. (e) Expression of p63 in the d0 p63GOF line, showing even with Dox treatment, there is no loss in Oct4 expression. (f) Strategy for generating the p63KO line, using a two gRNA CRISPR/Cas9 approach. (g) IF validation of the p63KO line, showing loss of p63 expression and higher levels of K18. All IF scale bars represent 50 μm. (h) Differential expression analysis from RNA-seq (measured by DESeq2) between the d0 and d0 p63GOF lines (upper panel), and the d7 p63WT and p63KO lines (lower panel). The gray dots on the scatter plot represent no change in gene expression between the two cell types, while red represents increased expression in the d0 or d7 wild type by a > 2 fold change and blue represents decreased expression in the d0 or d7 wild type by a < −2 fold change. Key transcription factors associated with epithelial development are induced by the morphogens and repressed by p63 at d7. (i) The p63 motif was the most significantly recovered motif under p63 ChIP-seq peaks in d0 p63GOF and d7 p63WT cells. (j) p63 binds distal to TSSs as depicted at the *HES1* locus (70 kb away) and to the same sites in d0 p63GOF and d7 p63WT. (k) p63 binds to similar sites genome-wide with and without morphogen presence, as depicted in the Venn Diagram. The majority of the d7 p63 binding sites are fully recovered in the d0 p63GOF line.

To assess the individual contributions to chromatin dynamics, we created p63 gain and loss of function hESCs using CRISPR/Cas9 technology (Fig. 1d,f) to yield a panel of four cell types: d0 (wild-type hESCs), d0 p63GOF (hESCs ectopically expressing p63), d7 p63WT (wild-type hESCs morphogen-treated, with endogenous p63), and d7 p63KO (hESCs morphogen-treated with no p63 expression). We verified p63 expression in these cell lines through IF, western blot, and sequencing (Fig. 1e,g, Extended Data Fig. 2).

Previous studies indicate that p63 overexpression can drive surface ectoderm commitment^21^, yet remarkably, expression of p63 in hESCs was insufficient to induce differentiation (Fig. 1e, Extended Data Fig. 2). Consistent with this observation, transcriptome analysis using RNA-sequencing (RNA-seq) revealed moderate changes in expression in roughly 300 genes between d0 and d0 p63GOF cells, whereas more than 2400 genes were differentially expressed in d7 p63WT vs. d7 p63KO cells (Fig. 1h).

Further, independent of the presence or absence of p63, morphogen exposure resulted in an exit from pluripotency and was required for p63 regulation of key transcription factors associated with epithelial development (Fig. 1h, Extended Data Fig. 2c). These important epithelial transcription factors, including TFAP2C, KLF4, GATA3, GRHL2, MSX2, and ELF3, were all repressed by p63 upon morphogen treatment. We conclude that morphogenetic signaling promotes a simple epithelial state, while enabling p63 to modify the morphogen-induced transcriptome to drive these stratified epidermal fates.

The striking influence of morphogens on p63 activity led us to investigate whether differences in p63 genomic occupancy accounted for the altered transcriptional activity. p63 ChIP-seq in d0 p63GOF and d7 p63WT revealed 7,960 and 6,097 p63 binding sites, respectively, and the p63 motif was significantly enriched under peaks in both datasets (Fig. 1i). Remarkably, over 70% of the sites were identical between both datasets (Fig. 1j,k), while 17% of peaks were gained in the d0 p63GOF (Extended Data Fig. 3a). Thus, differences in p63 occupancy cannot explain the dramatic morphogen-regulated p63 activity.

We next characterized how the morphogens and p63 affected chromatin accessibility and deposition of four histone modifications (H3K27ac, H3K27me3, H3K4me3, H3K4me1) using the Assay for Transposase Accessible Chromatin followed by sequencing (ATAC-seq) and histone chromatin immunoprecipitation (ChIP-seq), respectively. Overall, approximately 20,000 transposase accessible sites changed during the induction phase, with 14,000 opening and 6,000 closing between d0 and d7 p63WT (Fig. 2a). Additionally, over one third of the morphogen-dependent accessible sites became even more accessible upon p63 loss (Fig. 2a,d). Comparison of established histone modifications in d7 p63WT vs. p63KO revealed significant differences in H3K27me3, yet no observable differences on activating promoter or enhancer marks (Extended Data Fig. 3). Opposite to ATAC-seq changes, p63 absence resulted in a significant decrease in signal of the H3K27me3 mark, whereas H3K27me3 increased in d0 p63GOF cells (Fig. 2b,d). ChromHMM analysis indicated most of the accessibility changes and p63 binding sites occur in enhancers, rather than promoters (Extended Data Fig. 3). We conclude that p63 edits a subset of the morphogen-induced accessibility changes and regulates the accumulation of repressive H3K27me3 histone modifications.

**Figure 2.**
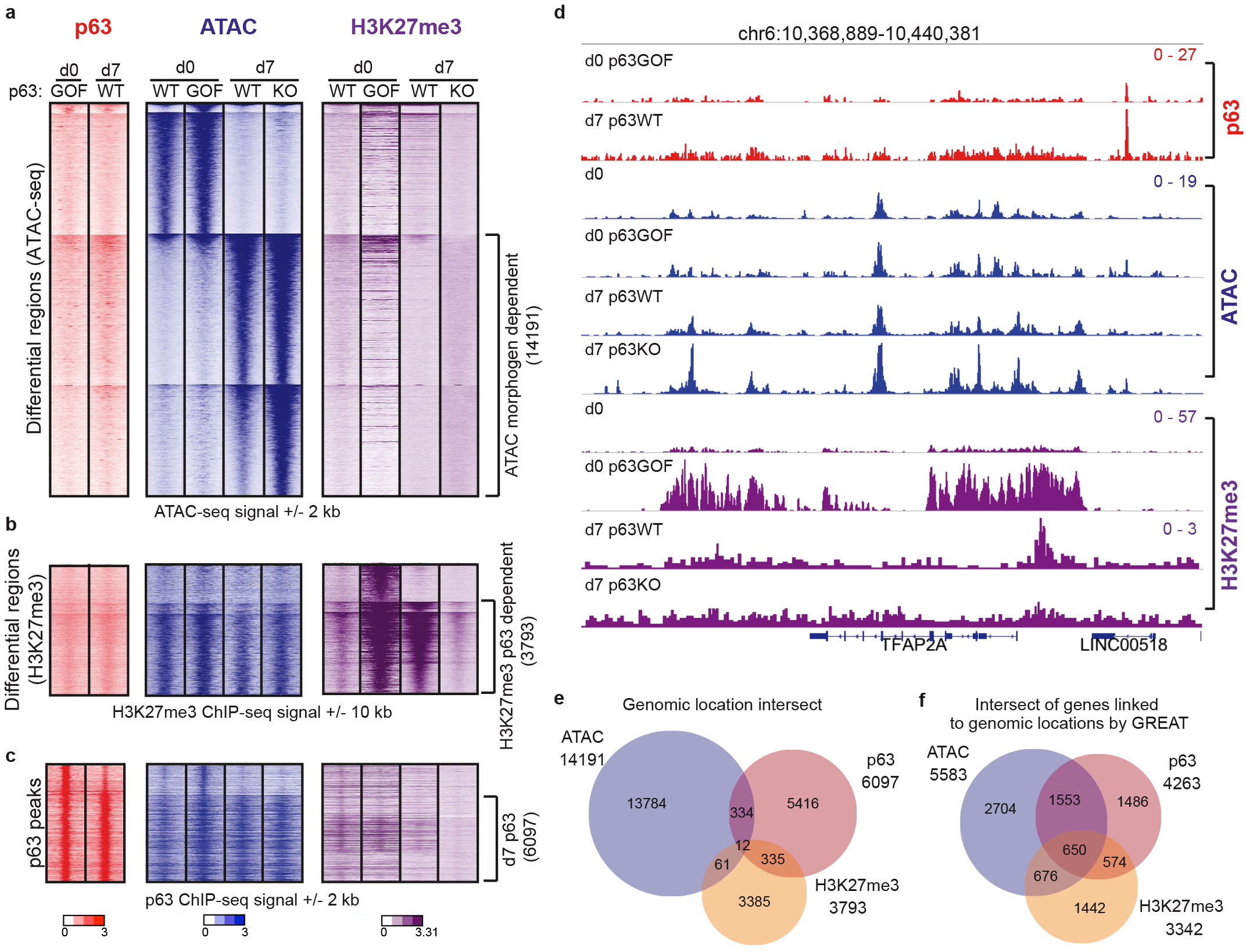
The morphogens establish an epigenetic landscape that p63 modifies at a distance. (a) Differential accessible regions between d0 and d7 p63WT as analyzed using DESeq2 on ATAC-seq signal. Heatmaps represent the signal at these ATAC regions within the various cell types and assays: p63 ChIP-seq signal (red, left panel), ATAC-seq signal (blue, middle panel), and H3K27me3 ChIP-seq signal (purple, right panel). 14,191 differential regions become more accessible upon morphogen treatment (morphogen-dependent). (b) Differential H3K27me3 regions between d7 p63WT and p63KO as analyzed by DESeq2. Heatmaps represent the same datasets as (a) only signal is shown at differential H3K27me3 sites (3,793 sites). (c) ATAC-seq (blue) and H3K27me3 (purple) signal at p63 binding sites (red). (d) Signal intensities of p63 ChIP-seq, ATAC-seq, and H3K27me3 ChIP-seq shown at the *TFAP2A* locus. (e) The overlap of genomic regions that are differential as measured in (a), (b), and (c). The genomic location intersect is very low. (f) GREAT analysis linking the above differential regions to the closest gene shows that these elements converge on a similar gene set. While their physical genomic locations do not overlap, they are linked to a common gene via GREAT.

Lineage selectors can act both directly on the epigenetic landscape at the site of binding to alter accessibility or histone modification deposition, or indirectly at a distance^22^. To determine how p63 acts, we intersected the p63-dependent H3K27me3 regions and morphogen-dependent accessible sites with p63 binding sites, revealing that few of the p63 binding sites overlapped with either of these changing elements (Fig. 2e). These data indicate that most of the p63 epigenetic regulatory action occurs distal to p63 binding. Interestingly, when we assigned p63 binding sites, morphogen-dependent accessible sites, and differential H3K27me3 regions to the nearest genes through GREAT, we found that these elements converge on a common gene set, despite each being distinct genomic regions (Fig. 2f, Extended Data Fig. 3).

To assess the connectivity and dynamics of the three-dimensional architecture between these distinct genomic regions, we employed cohesin HiChIP, a recent method analogous to Hi-C^10^, in all four cell types. We identified high-confidence chromatin contacts with 10 kb resolution using FitHiC^23^ (Extended Data Fig. 4) and demonstrated that 53% of p63 ChIP-seq peaks in d7 p63WT cells participate in these chromatin connections (Fig. 3a). Additionally, we illustrated that most morphogen and p63-dependent dynamic elements also participate in looping connections. Notably, only 34% of genes GREAT identified as having transcriptional start sites (TSSs) connected to p63 binding sites were verified by cohesin HiChIP, reinforcing the non-uniformity of the existing chromatin landscape (Extended Data Fig. 3,5).

**Figure 3.**
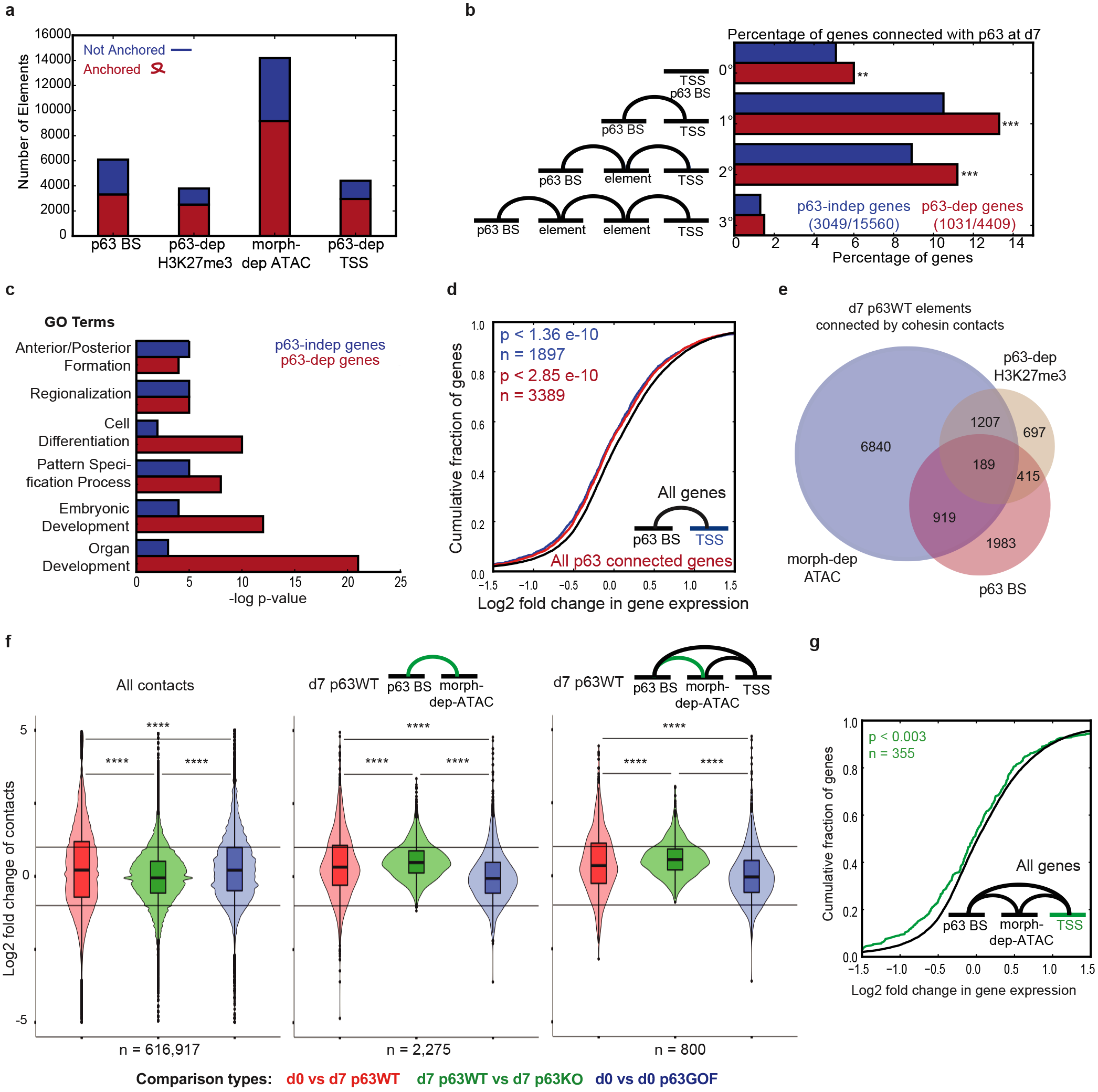
p63 - TSS connections are associated with negative regulation genome-wide. (a) Number of p63 binding sites (BS), p63-dependent (p63-dep) H3K27me3 sites, morphogen-dependent (morph-dep) ATAC sites, and p63-dependent TSSs that participate in chromatin looping (Anchored, red) vs those that do not (Not Anchored, blue) in d7 p63WT cells. (b) Percentage of p63-independent (p63-indep) genes (blue) and p63-dep genes (red), whose TSS is connected to p63 by direct binding (0°), direct contact (1°), or connected via one (2°) or two (3°) morph-dep ATAC and/or p63-dep H3K27me3 elements. (c) Gene Ontological Terms associated with p63-indep genes (blue) and p63-dep genes (red), which are connected to p63. (d) ecdf of the log2 fold change in gene expression between d7 p63WT vs d7 p63KO cells (d7 p63KO / d7 p63WT) for all p63 connected genes (red) and p63 1° connected genes (blue) compared to all genes (black). (e) 1° contact connections between p63 BS (red), p63-dep H3K27me3 (gold), and morph-dep ATAC (blue). (f) Change in connectivity strength between various cell types of all contacts (left panel), p63 - morph-dep ATAC contacts (middle panel), and p63 - morph-dep ATAC contacts in which both elements are connected to the TSS (right panel). n = number of contacts. (g) ecdf of the change in expression level (d7 p63WT vs d7 p63KO) of genes whose TSS is connected to a p63 BS and morph-dep ATAC site, which in turn are connected to each other (green) compared to all genes (black). FDR by monte carlo simulation **FDR<0.01,***FDR<0.001. Angela-Darling k-samples test ****p<1×10^−10^.

For the 4,409 protein-coding, p63-dependent genes, we determined the connectivity of their TSSs to a p63 binding site (Fig. 3b), revealing that 13% of these genes were in direct contact with p63 via chromatin looping (1º) and 11% were in contact via an indirect connection through a morphogen-dependent accessible site or H3K27me3 element (2º). Although more complex conformations through multiple elements (3º) were detected, random simulation demonstrated that p63 was not connected to p63-dependent genes by 3º connections at a frequency above random chance (FDR < 0.005); thus we focused on the 0º, 1 º, and 2º p63 connections (Fig. 3b, Extended Data Fig. 6).

We further interrogated the correlation between p63 connection to the TSS and transcriptional regulation, finding that p63 connectivity was insufficient to regulate gene expression. However, both p63-dependent and -independent genes connected to a p63 site were involved in organ development and cell differentiation, consistent with known p63 function (Fig. 3c)^7,8^. Additionally, the probability of transcriptional repression was significantly higher at genes connected to p63 (Fig. 3d). d7 p63-independent genes connected to p63 include keratinocyte differentiation genes whose expression becomes p63-dependent later during keratinocyte maturation, including p63 itself, MAFB, JAG1, ID2, and the Epidermal Differentiation Cluster (Extended Data Table 1)^24–26^. These data suggest that p63 and morphogen-regulated chromatin connections foreshadow future gene action. In all, we demonstrated that a large subset of the morphogen and p63-dependent elements are physically connected at d7 (Fig. 3e), accounting for the ability of p63 to regulate the epigenetic landscape at a distance.

Next, we determined the extent to which p63 and the morphogens influenced connectivity (Fig. 3f). In 1º (middle panel) and 2º (right panel) connections, contacts between morphogen-dependent accessible sites and p63 binding sites were regulated by both the morphogens and p63, with loss of p63 abolishing the connections, while overexpression of p63 failed to enhance them. Conversely, p63-H3K27me3 and p63-TSS interactions were enhanced by the morphogens and p63 overexpression, and weakened by p63 loss (Extended Data Fig. 7). Finally, we determined that of the 3D-conformations connecting p63 to a TSS, the connections to both morphogen-dependent accessible sites (Fig. 3g) and TSSs demonstrated greater repression than p63 connected via an H3K27me3 peak (Extended Data Fig. 7). These findings indicate that for optimal p63-regulated transcription both the morphogens and p63 are needed.

From our global analyses, we identified TFAP2C, a critical epithelial regulator^11^, as a gene induced by morphogens and repressed by p63 that exhibits a complex chromatin architecture driving its regulation. We sought to illustrate the p63-morphogen interactions by dissecting the p63 negative feedback regulation of this key developmental regulator (Fig. 4). Cohesin HiChIP and genomic analysis at this locus (Fig. 4a, Extended Data Fig. 8) revealed a distal p63 binding site with three d7 p63WT connections to the TSS: through a direct contact, the adjacent morphogen-dependent accessible site, and the distal H3K27me3 peak, all within 400 kb. We confirmed our cohesin HiChIP with UMI-4C^27^, a locus-specific technique, using primer viewpoints around the three connections (Extended Data Fig. 9).

**Figure 4.**
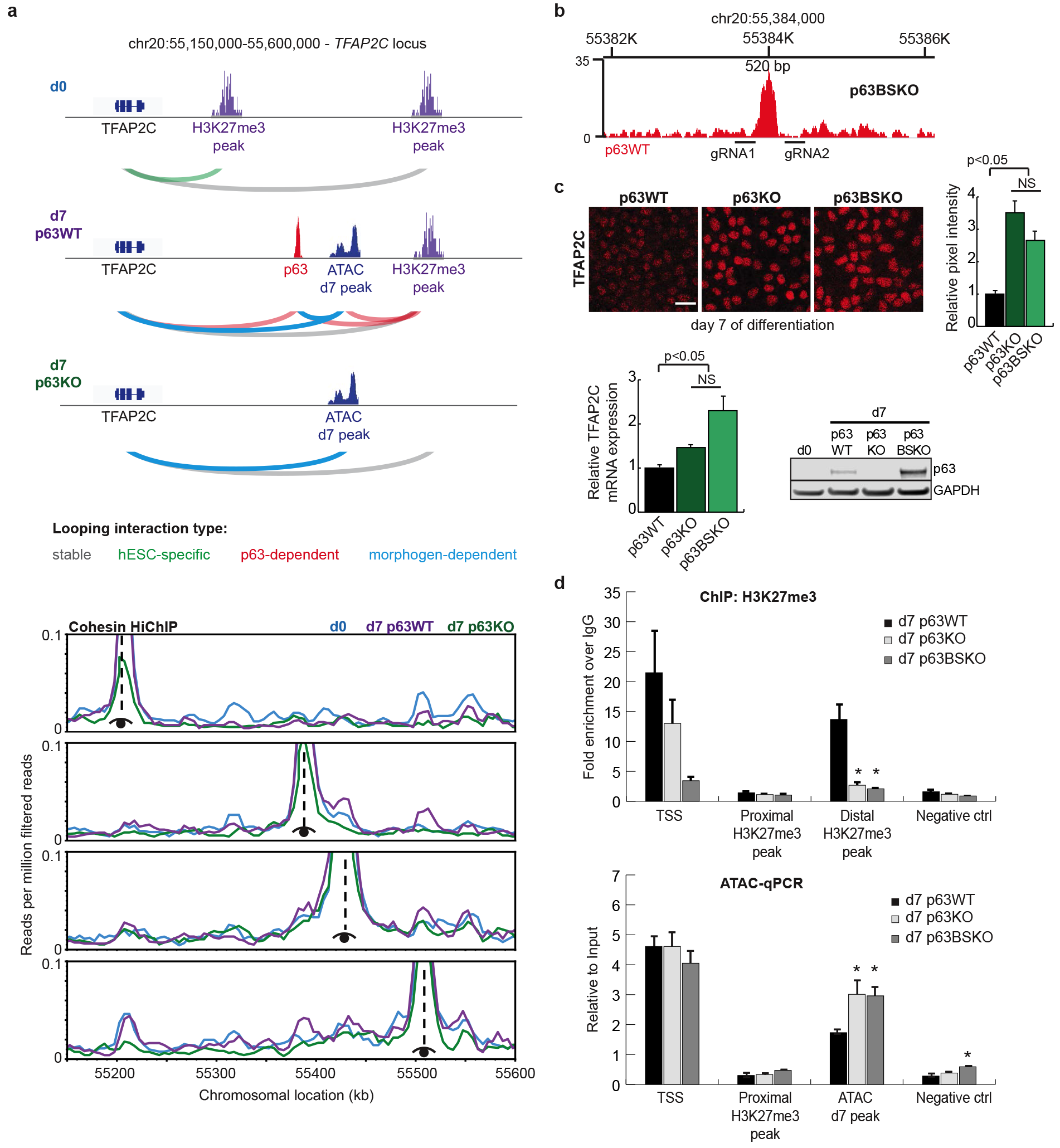
p63 negatively regulates TFAP2C expression through morphogen-induced and p63-dependent distal elements and connectivity. (a) Cohesin HiChIP reveals complex looping interactions between numerous distal elements at the *TFAP2C* locus. Schematic for the morphogen and p63-dependent interactions (top panel) with virtual 4C plots of the normalized cohesin HiChIP data below (bottom panel). (b) The p63 binding site was deleted using CRISPR/Cas9 to determine the effects of its loss on TFAP2C expression. p63BSKO is a 520 bp deletion surrounding the p63 ChIP-seq peak. (c) Deletion of the p63 binding site leads to an increase in TFAP2C expression similar to the levels seen in the d7 p63KO cells (NS - not significant). Loss of TFAP2C expression also leads to a dramatic increase in p63 expression. Relative pixel intensity was calculated from 3 independent images and scale bars represent 20 μm. (d) ChIP-qPCR for H3K27me3 at the *TFAP2C* locus shows a decrease in the histone mark in the d7 p63BSKO cells, similar to the d7 p63KO cells (*p-value < 0.01). ATAC-qPCR at this locus shows an increase in accessibility at the d7 ATAC peak in d7 p63BSKO, again similar to the d7 p63KO (*p-value < 0.05). Deletion of the p63 binding site results in a loss of tightly controlled TFAP2C expression. Both graphs depict signal relative to input and error bars represent standard error of the mean.

Comparison of the chromosome conformation among the different cell lines indicated that p63 presence enhances connectivity to all three of the main loops at d7 and in the absence of p63, the connections and transcriptional output collapse. Morphogen exposure connects p63 to the induced neighboring morphogen-dependent accessible site, but the connection relies on ongoing p63 expression to maintain it, as loss of p63 fails to uphold it despite morphogen presence. Thus, our analysis of the *TFAP2C* locus shows that both the morphogens and p63 contribute to proper regulation.

To validate the importance of the morphogen-dependent accessible site, we removed the region using CRISPR/Cas9 and demonstrated a loss of morphogen-induced TFAP2C expression (Extended Data Fig. 8b). Furthermore, we hypothesized that removal of the p63 binding site should drive both TFAP2C and p63 expression, given our observation that TFAP2C induces p63 expression in hESCs (unpublished results) and that p63 provides important early negative regulation of TFAP2C. To test this, we deleted the p63 binding site (p63BSKO) and found dramatically elevated levels of TFAP2C at d7, consistent with the predicted negative feedback modulation of TFAP2C by p63 (Fig. 4b,c). Moreover, d7 p63BSKO cells showed increased expression of p63, demonstrating the need for tight p63-morphogen regulation to control the levels of key developmental factors. Histone ChIP-qPCR revealed a loss of H3K27me3 accumulation at both the TSS and the distal H3K27me3 site in d7 p63BSKO cells, while other non p63-connected sites remained unaffected (Fig. 4d). Similarly, the morphogen-dependent accessible site became more accessible in d7 p63BSKO cells, to levels found in d7 p63KO cells (Fig. 4d), confirming the connectivity of these distal elements.

Here we deepen our understanding of the interplay between morphogens and lineage selectors during surface ectoderm commitment, and find the surprising inability for the lineage selector p63 to function in the absence of morphogen action. Morphogens provide the powerful driving force for cell state change by inducing expression of the lineage factor while also altering chromatin accessibility, histone modifications, and chromosome conformation. p63, in turn, further modifies the morphogen-dependent epigenetic landscape to drive surface ectoderm differentiation. Further, our results illustrate how chromatin connections to the lineage selector p63 are necessary and more likely to induce gene expression changes, but are not sufficient. Our finding that p63 at d7 is poised to act on later keratinocyte differentiation genes (Extended Data Table 1)^24–26^ suggests the existence of additional inductive influences after addition of RA/BMP that will enable broader p63-dependent transcription. This is functionally similar to “poised” histone modifications and provides a structural explanation of how the order of morphogen exposure can determine downstream transcriptional programs. This study has important implications for the apparent autonomy of lineage selectors and for the basis of morphogenesis. Our work suggests that small changes in morphogen activity can dramatically alter the induced chromosome landscape and connectivity, explaining how a single lineage selector like p63 can direct such a panoply of transcriptional programs depending on the specific morphogen exposure.

## Acknowledgements

We thank members of the Oro Laboratory, P. Greenside, J. Wysocka, A. Kundaje, O. Wapinski, and D. Webster for helpful discussions and comments. This work was supported by CIRM Tools grant RT3-07796 (A.E.O.), NIH/NIAMS grant F32AR070565 (J.M.P.), and NIH P50 HG007735 (H.Y.C.).

## Author Contributions

S.P.M. and J.M.P. designed and executed experiments, analyzed data, and wrote the manuscript. S.N.P. analyzed data and edited the manuscript. J.L.T., E.B., M.R.M., C.R., H.H.Z., and L.L. executed experiments and contributed to experimental design. E.L., D.A., A.J.R., and G.S. contributed to data analysis. H.Y.C. and P.A.K. contributed to experimental design. A.E.O. designed experiments, analyzed data, wrote the manuscript, and conceived the project with S.P.M.

## Methods

### CRISPR/Cas9 guided genome editing

gRNAs were designed using the online tool available at http://crispr.mit.edu/^28^, selected based on the highest scores and the least off-targets, and incorporated into a DNA fragment bearing all the components necessary for gRNA expression^29^. Donor sequences were designed by selecting 700 bp arms flanking left and right of the region to be modified. Both gRNAs and donor sequences were synthesized as 5-phosphorylated gene blocks (IDT) and cloned into a blunted plasmid with puromycin selection, except for gRNAs targeting the AAVS1 locus, which were acquired through Addgene (Plasmid # 72833)^30^. The d0 p63GOF line was generated by integrating the humanized p63 mouse cDNA under the control of a Tetracycline Responsive Element (TRE) to the AAVS1 locus. Doxycycline (Sigma) was added to the media for 2 days at a concentration of 2 ug/ml to induce expression of p63 in hESCs.

### hESC culture and transfection

H9 human embryonic stem cells were cultured on Vitronectin Recombinant Human Protein (Life Technologies) in Essential 8 medium (E8, Life Technologies) as described previously^16^. Cells were passaged every three days as clumps with 0.5 mM EDTA (Lonza). For transfection, 2×10^6^ cells were nucleofected using AmaxaTM P3 Primary Cell 4D-Nucleofector (Lonza) as recommended by the manufacturer, with no more than a 10 uL mix of 2 ug of plasmid carrying each gRNA, 2 ug of plasmid carrying hCas9 and 2-4 ug of plasmid carrying the donor DNA to repair the Cas9/gRNA induced break by homologous recombination. Cells were plated and allowed to recover for a minimum of 6h in E8 media supplemented with 2 uM thiazovivin (Stemgent). Drug selection with 1 ug/mL puromycin (InvivoGen) started 48h after transfection and lasted 2 days for loss of function cell lines, or continued for several days for gain of function cell line. Colonies were picked 10 days after selection and genotyped by PCR to confirm homozygosity.

### *In vitro* epithelial hESC differentiation

For differentiation, 6.2×10^3^ cells/cm^2^ were plated as colonies on Vitronectin coated plates. Next day, media was changed to Essential 6 (E6, Life Technologies) supplemented with 1 uM RA (Sigma) and 5 ng/mL Recombinant Human BMP-4 (R&D Systems), and replaced every two days for seven days, at which point cells were dissociated with Accutase (StemCell Technologies) and collected for downstream analysis, or media was replaced to Defined Keratinocyte-SFM media (DKSFM, Life Technologies) for terminal differentiation into keratinocytes.

### Immunofluorescence staining

Cells were cultured on glass cover slips in 12 wells, subjected to the appropriate treatment and fixed for 10 min at room temperature in 4% paraformaldehyde in PBS. Cells were permeabilized for 10 min with permeabilization buffer (0.1% Triton-X + 0.05% Tween-20 in PBS) and blocked for 30 min with 10% Horse Serum (Vector Laboratories) in permeabilization buffer. Antibodies at appropriate dilutions were incubated overnight at 4°C. Secondary antibodies were added at 1:500 dilution and incubated at room temperature protected from light for 1h. Cells were washed three times in Hoechst 1:10,000 in PBS, and glass cover slips were mounted onto glass slides with mounting medium before imaging. Antibodies were diluted in permeabilization buffer at the indicated dilutions: AP-2γ (1:100, Cell Signaling 2320S), p63 (1:200, Genetex GTX102425), KRT18 (1:800, R&D AF7619), KRT14 (1:1000, BioLegend 906001), OCT4 (1: 100, BioLegend 631902).

### RNA extraction and library preparation

For RNA extraction, cells were lysed directly in Trizol (Invitrogen), purified as indicated by the manufacturer, and then run through RNeasy columns (Qiagen). Libraries for RNA-seq were prepared using TrueSeq RNA Library Prep Kit v2 (Illumina) according to the manufacturer’s protocol. Real time PCR was performed with SYBR Green PCR master mix (Life Technologies) and in a Stratagene real time PCR machine.

### Chromatin immunoprecipitation (ChIP) and library preparation

Cells were cross-linked in suspension for 10 min using freshly prepared 1% formaldehyde (Thermo Scientific) in PBS. Subsequently, glycine was added to a final concentration of 0.125 M to quench formaldehyde, and cells were washed twice with cold PBS. 60×10^6^ or 10×10^6^ cross-linked cells were used per ChIP for p63 or histone marks, respectively. Cells were lysed in lysis buffer (50 mM Tris-HCl pH 8.0, 10 mM EDTA, 0.5% SDS, 1X protease inhibitors) for 30 minutes on ice and sonicated for 2h using a Bioruptor (Diagenode) to achieve a chromatin size between 200 and 300 bp. Chromatin was centrifuged to remove debris, quantified and diluted in dilution buffer (50 mM Tris-HCl pH 8.0, 10 mM EDTA, 1X protease inhibitors) to achieve a 0.1% SDS final concentration. Sheared chromatin was incubated overnight at 4° with appropriate antibodies, followed by incubation with 30 uL of agarose G beads (Invitrogen) for 4h at 4°C. Antibodies were used at the indicated concentrations per ChIP (per 10×10^6^ cells): p63 (12 uL, Active Motif 39739), H3K4me3 (5 ug, Abcam ab8580), H3K4me1 (5 ug, Abcam ab8895), H3K27Ac (5 ug, Abcam ab4729), H3K27me3 (5 ug, Millipore 07-449). Beads were washed twice each with low salt buffer (50 mM Tris-HCl pH 8.0, 0.15 M NaCl, 1 mM EDTA pH 8.0, 0.1% SDS, 1% triton X-100, 0.1% sodium deoxycholate), high salt buffer (50 mM Tris-HCl pH 8.0, 0.5 M NaCl, 1 mM EDTA pH 8.0, 0.1% SDS, 1% triton X-100, 0.1% sodium deoxycholate), and LiCl buffer (50 mM Tris-HCl pH 8.0, 0.15 M LiCl, 1 mM EDTA pH 8.0, 1% Nonidet P-40, 0.1% sodium deoxycholate). DNA was eluted in 100 uL of elution buffer (50 mM NaHCO_3_, 1% SDS) and crosslinks were reversed with 4 uL of 5 M NaCl incubated overnight at 67°C. RNA was removed by adding 1 uL of 10 mg/mL RNase A and incubating for 30 min at 37°C. DNA was cleaned using the Qiagen Qiaquick PCR purification kit and quantified using Qubit (Invitrogen). Between 5 and 10 ng of pooled DNA were used for library preparation using NEBNext kit (New England Biolabs) and Agencourt AMPure XP beads (Beckman) according to the manufacturer’s protocol. Single-read libraries were sequenced on Illumina NextSeq sequencer.

### Assay for Transposase Accessible Chromatin (ATAC-seq)

ATAC-seq was performed as described^31^. Briefly, after treatment with Accutase, 7×10^4^ cells were washed with cold PBS and lysed using 0.1% NP40 in RSB buffer. Nuclei pellets were Tn5 transposed using the DNA Sample Preparation Kit from Nextera®. Libraries were amplified for 9-15 total cycles using the Ad1 and Ad 2.1-2.16 barcodes. Libraries were purified using the Min-Elute columns (Qiagen) and eluted with 10 μL of buffer EB. Library DNA concentrations were determined with Bioanalyzer High-Sensitivity DNA analysis kit (Agilent). Paired-end libraries were sequenced initially on a MiSeq sequencer and analyzed using a custom script to determine the signal enrichment over background at TSSs over a 2 kb window. (https://www.encodeproject.org/data-standards/atac-seq/) Only libraries that had enrichment scores above 6 were sequenced deeper in a NextSeq Illumina sequencer.

### Cohesin HiChIP

In situ chromosome conformation capture (3C) was performed as described earlier^10^. Briefly, 25×10^6^ cells were crosslinked and digested with MboI (NEB). After digest, biotin was incorporated into the sticky ends of fragments before ligation. Cohesin ChIP was performed to enrich for proximity ligations bound to cohesin, using an SMC1 antibody (Bethyl, A300-055A). The library quality was assessed on a MiSeq sequencer before sequencing on an Illumina HiSeq. Three replicates were pooled and sequenced across two HiSeq lanes for a total of 1200 million reads per sample.

### UMI-4C

UMI-4C was performed as described previously^27^. Briefly, 1×10^7^ cells were crosslinked in suspension with 1% formaldehyde then quenched with glycine, and pelleted cells were lysed in 1 mL fresh cold lysis buffer (50 mM Tris-HCl, pH 7.5, 150 mM NaCl, 5 mM EDTA, 0.5% NP-40, 1% TX-100, 1x protease inhibitors) on ice. The nuclei were extracted and resuspended in water, DpnII buffer, and 10% SDS for DpnII digestion. Three rounds of DpnII digestion were performed, adding 200 U of HC DpnII (NEB) for 2 hours, incubating overnight, and then 2 more hours all at 37°C with rotation. After inactivation of DpnII, the 3C reactions were diluted to 7 mL and 13.6 uL of HC T4 ligase (NEB) were added for overnight ligation at 16°C. Crosslinks were reversed overnight at 65°C with Proteinase K (Qiagen) and DNA was treated with RNase A (Qiagen) for 45 minutes at 37°C. The DNA was then purified with one phenol-chloroform extraction (ThermoScientific) and ethanol precipitation, and resuspended in 150 uL of 10 mM Tris-HCl, pH 8.0. DNA was quantified using Qubit before proceeding with library preparations. Aliquots of chromatin were taken before and after DpnII digestion and after overnight ligation to determine efficiency of enzymatic reactions. UMI-4C library preparation was performed as described previoiusly^27^. Briefly, 5-10 ug of 3C library was sonicated in a Diagenode Bioruptor to achieve a chromatin size between 400 and 600 bp. End-repair (NEB kit), A-tailing (NEB kit) and 5’-dephosphorylation (NEB) of ends were performed as recommended by the manufacturer. TruSeq Illumina indexed adapters were ligated to the 3’-end of the DNA using Quick Ligase (NEB). Libraries were generated by nested PCR at particular genomic loci using GoTaq Hot Start Polymerase (Promega) and 200 ng of DNA template (Extended Data Fig. 9). The primer for the second PCR included the Illumina dangling adaptor for enrichment of the product from the first PCR, as described^27^. Paired-end libraries were sequenced in the NextSeq sequencer.

### Next Generation Sequencing processing of ChIP-seq and ATAC-seq data

Quality control of fastq files was done with FASTQC^32^. Sequence alignment to hg19 was performed using tophat for RNA-seq (parameters: p 10 --library-type fr-firststrand -r 100 --mate-std-dev 100), or bowtie2^33^ for ChIP-seq (parameters: -p 24 -S -a -m 1 --best – strata) and ATAC-seq (parameters: -p 24 -S -m 1 -X 2000). Aligned reads were processed to remove PCR duplicates using samtools^34^ and mitochondrial DNA (for ATAC-seq datasets). Peak calling was carried out with MACS2^35^ using default settings with a p-value of 0.05. To filter out non-reproducible peaks, called peaks from biological replicates were processed through the Irreproducible Discovery Rate (IDR) framework implemented in R^36^.

### Differential counting, heatmaps, and average profiles

For ChIP-seq and ATAC-seq, a union list of the MACS2 called peaks per sample not filtered by IDR was generated using bedtools merge^37^, and raw reads covering each region were recovered from bam files using bedtools multicoverage. For RNA-seq, raw reads on reference genes were recovered using HOMER (version 4.8^38^ analyzeRepeats.pl command). To test for differential counting, raw reads were compared using DESeq2 package implemented in R^39^, and filtered based on an adjusted p-value of < 0.05 and 2 fold change. For heatmaps and average profiles, tag counts were recovered +/− 2 kb from the peak summit using HOMER annotatePeaks.pl command with -hist 25 -ghist or -hist 25 parameters. Heatmap images were generated using Java TreeView^40^. Average profiles and scatter plots were plotted using Python matplotlib.

### Motif discovery and Gene Ontology

*De novo* motif discovery was performed using Homer findMotifsGenome function with – size 200 as a parameter. Gene ontology analysis was performed using DAVID^41^ for RNA-seq data and GREAT^42^ for ChIP-seq and ATAC-seq data.

### Chromatin state determination

ChromHMM software^43^ was used to learn and identify chromatin states as instructed in the manual. Encode chromatin segmentation using ChromHMM was used as a reference to label each state using a custom script using bedtools intersect. The enrichment of each state for a set of peaks was calculated using the NeighborhoodEnrichment command and compared among samples using a custom script. Enrichments were plotted using Python matplotlib library.

### Analysis of UMI-4C data

UMI-4C data was aligned and analyzed using HiC-Pro^44^ and the DpnII segmented genome annotation file. Interaction matrices of 5 kb resolution were generated and used to create Virtual 4C profiles through a custom python script and the matplotlib library.

### Analysis of cohesin HiChIP data

HiChIP paired end reads were aligned to hg19 using HiC-Pro^44^. Duplicate reads were removed, assigned to MboI restriction fragments, filtered for valid interactions, and then used to generate binned interaction matrices of both 5 kb and 10 kb resolution. The 5 kb interaction matrices were used to visualize contacts by Virtual 4C, similar to the UMI-4C analysis. The 10 kb interaction matrix was used to call high confidence contacts (defined as counts ≥ 10, FDR < 0.001) using the contact caller, FitHiC^23^. Default FitHiC settings were used to generate an FDR for each bin pair. These high confidence cohesin contacts were used in the subsequent analyses.

### Contact connection analyses

An element was considered participating, or anchored, in cohesin connections, if it possessed at least one high confidence contact bin in a given cell type. When considering ways in which p63 was connected to a TSS (defined as TSS +/− 5 kb), four chromatin conformations were considered. 0° connections were defined as two elements overlapping in physical space (e.g. p63 BS contained within the TSS). 1° connections were defined as one element anchored in one bin of a cohesin contact and the second element anchored in the other bin. More complicated connections between elements were also considered: 2° connections were defined as two elements in distinct physical space both forming 1° connections to the same third element. Finally, 3° connections were when one element formed a 1° connection to a second element, which also formed a 1° connection to a third element, which also had a 1° connection to the fourth (target) element. All elements in both 2° and 3° configurations were in distinct physical space (i.e. non-overlapping).

### Differential contact analysis

The Bioconductor package edgeR^45^ was used to perform multiple comparison differential analysis of high confidence FitHiC contacts in d0, d0 p63GOF, d7 p63WT, and d7 p63KO cells. The Anderson-Darling K-sample test, a modification of the K-S test, which gives greater weight to the tails, was used to calculate statistical significance between populations of the fold change in contact connectivity^46^.

### Code Availability

Custom scripts described in the Methods will be made available upon request.

### Data Availability

All sequencing data will be available through Gene Expression Omnibus (GEO) - accession number pending.

A **Life Sciences Reporting Summary** for this publication is available.

## References

1. Walmsley, G.G., et al. Induced pluripotent stem cells in regenerative medicine and disease modeling. Curr Stem Cell Res Ther 9, 73–81 (2014).

2. Inoue, H., Nagata, N., Kurokawa, H. & Yamanaka, S. iPS cells: a game changer for future medicine. The EMBO journal 33, 409–417 (2014).

3. Hanna, J., et al. Treatment of sickle cell anemia mouse model with iPS cells generated from autologous skin. Science 318, 1920–1923 (2007).

4. Umegaki-Arao, N., et al. Induced pluripotent stem cells from human revertant keratinocytes for the treatment of epidermolysis bullosa. Sci Transl Med 6, 264ra164 (2014).

5. Sebastiano, V., et al. Human COL7A1-corrected induced pluripotent stem cells for the treatment of recessive dystrophic epidermolysis bullosa. Sci Transl Med 6, 264ra163 (2014).

6. Zaret, K.S. & Carroll, J.S. Pioneer transcription factors: establishing competence for gene expression. Genes & development 25, 2227–2241 (2011).

7. Yang, A., et al. p63 is essential for regenerative proliferation in limb, craniofacial and epithelial development. Nature 398, 714–718 (1999).

8. Mills, A.A., et al. p63 is a p53 homologue required for limb and epidermal morphogenesis. Nature 398, 708–713 (1999).

9. Lupien, M., et al. FoxA1 translates epigenetic signatures into enhancer-driven lineage-specific transcription. Cell 132, 958–970 (2008).

10. Mumbach, M.R., et al. HiChIP: efficient and sensitive analysis of protein-directed genome architecture. Nature methods 13, 919–922 (2016).

11. Qiao, Y., et al. AP2gamma regulates neural and epidermal development downstream of the BMP pathway at early stages of ectodermal patterning. Cell research 22, 1546–1561 (2012).

12. Metallo, C.M., Ji, L., de Pablo, J.J. & Palecek, S.P. Retinoic acid and bone morphogenetic protein signaling synergize to efficiently direct epithelial differentiation of human embryonic stem cells. Stem cells 26, 372–380 (2008).

13. Itoh, M., et al. Generation of 3D skin equivalents fully reconstituted from human induced pluripotent stem cells (iPSCs). PloS one 8, e77673 (2013).

14. Guenou, H., et al. Human embryonic stem-cell derivatives for full reconstruction of the pluristratified epidermis: a preclinical study. Lancet 374, 1745–1753 (2009).

15. Coraux, C., et al. Reconstituted skin from murine embryonic stem cells. Curr Biol 13, 849–853 (2003).

16. Chen, G., et al. Chemically defined conditions for human iPSC derivation and culture. Nature methods 8, 424–429 (2011).

17. Owens, D.W. & Lane, E.B. The quest for the function of simple epithelial keratins. BioEssays: news and reviews in molecular, cellular and developmental biology 25, 748–758 (2003).

18. Senoo, M., Pinto, F., Crum, C.P. & McKeon, F. p63 Is essential for the proliferative potential of stem cells in stratified epithelia. Cell 129, 523–536 (2007).

19. Koster, M.I. & Roop, D.R. Mechanisms regulating epithelial stratification. Annu Rev Cell Dev Biol 23, 93–113 (2007).

20. Green, H., Easley, K. & Iuchi, S. Marker succession during the development of keratinocytes from cultured human embryonic stem cells. Proceedings of the National Academy of Sciences of the United States of America 100, 15625–15630 (2003).

21. Aberdam, E., et al. A pure population of ectodermal cells derived from human embryonic stem cells. Stem cells 26, 440–444 (2008).

22. Wamstad, J.A., et al. Dynamic and coordinated epigenetic regulation of developmental transitions in the cardiac lineage. Cell 151, 206–220 (2012).

23. Ay, F., Bailey, T.L. & Noble, W.S. Statistical confidence estimation for Hi-C data reveals regulatory chromatin contacts. Genome research 24, 999–1011 (2014).

24. Barton, C.E., et al. Novel p63 target genes involved in paracrine signaling and keratinocyte differentiation. Cell death & disease 1, e74 (2010).

25. Koh, L.F., Ng, B.K., Bertrand, J. & Thierry, F. Transcriptional control of late differentiation in human keratinocytes by TAp63 and Notch. Exp Dermatol 24, 754–760 (2015).

26. Truong, A.B., Kretz, M., Ridky, T.W., Kimmel, R. & Khavari, P.A. p63 regulates proliferation and differentiation of developmentally mature keratinocytes. Genes & development 20, 3185–3197 (2006).

27. Schwartzman, O., et al. UMI-4C for quantitative and targeted chromosomal contact profiling. Nature methods 13, 685–691 (2016).

## References

28. Hsu, P.D., et al. DNA targeting specificity of RNA-guided Cas9 nucleases. Nature biotechnology 31, 827–832 (2013).

29. Mali, P., Esvelt, K.M. & Church, G.M. Cas9 as a versatile tool for engineering biology. Nature methods 10, 957–963 (2013).

30. Natsume, T., Kiyomitsu, T., Saga, Y. & Kanemaki, M.T. Rapid Protein Depletion in Human Cells by Auxin-Inducible Degron Tagging with Short Homology Donors. Cell Rep 15, 210–218 (2016).

31. Buenrostro, J.D., Giresi, P.G., Zaba, L.C., Chang, H.Y. & Greenleaf, W.J. Transposition of native chromatin for fast and sensitive epigenomic profiling of open chromatin, DNA-binding proteins and nucleosome position. Nature methods 10, 1213–1218 (2013).

32. Andrews, S. FastQC: a quality control tool for high throughput sequence data. . (2010).

33. Hwang, S., Kim, E., Lee, I. & Marcotte, E.M. Systematic comparison of variant calling pipelines using gold standard personal exome variants. Sci Rep 5, 17875 (2015).

34. Li, H., et al. The Sequence Alignment/Map format and SAMtools. Bioinformatics 25, 2078–2079 (2009).

35. Zhang, Y., et al. Model-based analysis of ChIP-Seq (MACS). Genome biology 9, R137 (2008).

36. Li, H. A statistical framework for SNP calling, mutation discovery, association mapping and population genetical parameter estimation from sequencing data. Bioinformatics 27, 2987–2993 (2011).

37. Quinlan, A.R. & Hall, I.M. BEDTools: a flexible suite of utilities for comparing genomic features. Bioinformatics 26, 841–842 (2010).

38. Heinz, S., et al. Simple combinations of lineage-determining transcription factors prime cis-regulatory elements required for macrophage and B cell identities. Molecular cell 38, 576–589 (2010).

39. Love, M.I., Huber, W. & Anders, S. Moderated estimation of fold change and dispersion for RNA-seq data with DESeq2. Genome biology 15, 550 (2014).

40. Saldanha, A.J. Java Treeview--extensible visualization of microarray data. Bioinformatics 20, 3246–3248 (2004).

41. Huang da, W., Sherman, B.T. & Lempicki, R.A. Systematic and integrative analysis of large gene lists using DAVID bioinformatics resources. Nature protocols 4, 44–57 (2009).

42. McLean, C.Y., et al. GREAT improves functional interpretation of cis-regulatory regions. Nature biotechnology 28, 495–501 (2010).

43. Ernst, J. & Kellis, M. ChromHMM: automating chromatin-state discovery and characterization. Nature methods 9, 215–216 (2012).

44. Servant, N., et al. HiC-Pro: an optimized and flexible pipeline for Hi-C data processing. Genome biology 16, 259 (2015).

45. Anders, S., et al. Count-based differential expression analysis of RNA sequencing data using R and Bioconductor. Nature protocols 8, 1765–1786 (2013).

46. Scholz, F. & Stephens, M. K-Sample Anderson-Darling Tests. J Am Stats Assoc 82, 918–924 (1987).

